# Improvement of phycocyanobilin synthesis for genetically encoded phytochrome-based optogenetics

**DOI:** 10.1101/2020.01.15.908343

**Authors:** Youichi Uda, Haruko Miura, Yuhei Goto, Kazuhiro Aoki

## Abstract

Optogenetics is a powerful technique using photoresponsive proteins, and light-inducible dimerization (LID) systems, an optogenetic tool, allow to manipulate intracellular signaling pathways. One of the red/far-red responsive LID system, phytochrome B (PhyB)-phytochrome interacting factor (PIF), has a unique property of controlling both association and dissociation by light on the second time scale, but PhyB requires a linear tetrapyrrole chromophore such as phytochromobilin or phycocyanobilin (PCB), and such chromophores are present only in higher plants and cyanobacteria. Here, we report that we further improved our previously developed PCB synthesis system (synPCB), and successfully established a stable cell line containing a genetically encoded PhyB-PIF LID system. First, four genes responsible for PCB synthesis, namely, *PcyA*, *HO1*, *Fd*, and *Fnr*, were replaced with their counterparts derived from thermophilic cyanobacteria. Second, Fnr was truncated, followed by fusion with Fd to generate a chimeric protein, tFnr-Fd. Third, these genes were concatenated with P2A peptide cDNAs for polycistronic expression, resulting in an approximately 4-fold increase in PCB synthesis compared with the previous version. Finally, we incorporated PhyB-PIF and synPCB into drug inducible lentiviral and transposon vectors, which enabled us to induce PCB synthesis and PhyB-PIF LID system by doxycycline treatment. These tools provide a new opportunity to advance our understanding of the causal relationship between intracellular signaling and cellular functions.

## Introduction

Optogenetics is an emerging technique that employs photoreceptor proteins to control protein and cellular functions by light ^1–3^. Among optogenetic tools, light-induced dimerization (LID) systems allow the optical manipulation of intracellular signaling ^4–7^, gene expression ^8^ and organelle transport ^9^. The *Avena sativa* light-oxygen-voltage (LOV) domain and *Arabidopsis thaliana* Cryptochrome 2 (CRY2) have been widely used as photoresponsive proteins for light-induced protein-protein hetero-dimerization ^10,11^. This is partly because LOV and CRY2 do not require an exogenous chromophore such as flavin mononucleotide (FMN) or flavin adenine dinucleotide (FAD). Both FMN and FAD are abundant in many organisms, and therefore LOV- or CRY-based LID system can be achieved merely by expressing these proteins. Meanwhile, LOV and CRY2 exert their optogenetic effects in response to blue light, making it difficult to be used together with fluorescent proteins such as CFP and GFP. In addition, blue light illumination causes phototoxicity to many live organisms.

*Arabidopsis thaliana* phytochrome B (PhyB), which is a red/far-red light-absorbing photoreceptor protein involved in the control of developmental responses ^12,13^, has been utilized as an optogenetic tool to manipulate gene expression and intracellular signaling ^14–16^. PhyB requires a chromophore such as phytochromobilin (PΦB) or phycocyanobilin (PCB) for the photoreception ^17^. The chromophore covalently attaches to apo-PhyB, generating holo-PhyB ^18^. The holo-PhyB protein acts as a reversible switch in light perception; it provides photointerconversion between the red light-absorbing ground state of PhyB (Pr) and far-red light-absorbing active state of PhyB (Pfr). The PhyB (Pfr) associates with effector proteins such as phytochrome interacting factors (PIFs)^19^(Figure 1A). The PhyB-PIF LID system has a unique feature, showing rapid association and dissociation kinetics induced by red- and far-red-light, respectively. However, if the PhyB system is used in organisms other than photosynthetic organisms, chromophores, namely, PΦB and PCB, are required. Although researchers have added PCB purified from algae to function optogenetics with PhyB, the resulting PhyB system is technically difficult to apply *in vivo*, where delivery and clearance may be limitted.

**Figure 1.**
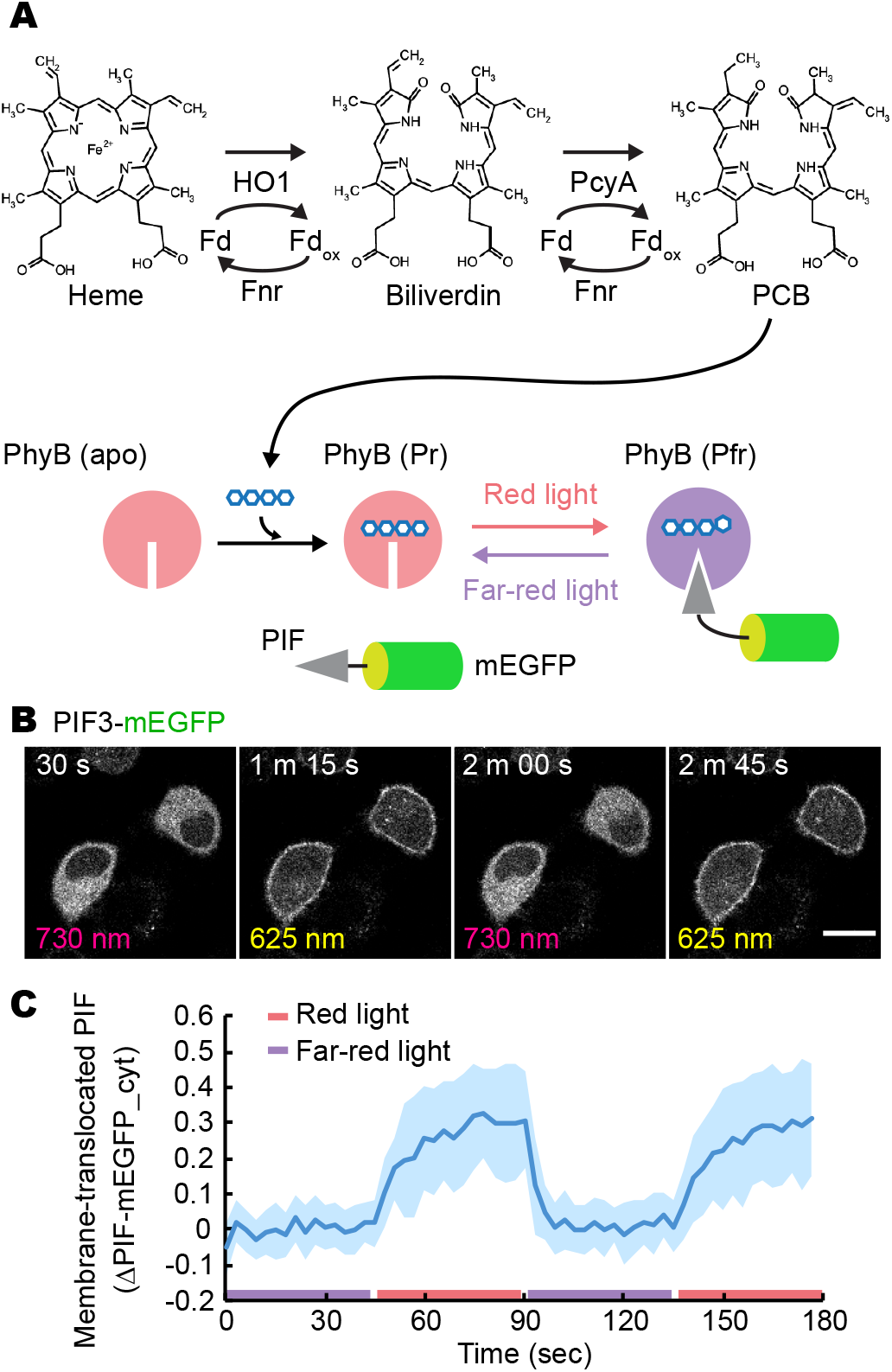
PhyB-PIF optogenetic switch with PCB synthesis in mammalian cells. (A) The metabolic pathway for PCB synthesis. First, heme is catalyzed by HO1 with Fd and Fnr to generate biliverdin. Then, PcyA with Fd and Fnr produces PCB, which is covalently attached to apo-PhyB. Upon activation by red light irradiation, the red light-absorbing form of PhyB (Pr) is converted to the far-red light-absorbing form of PhyB (Pfr), which binds to PIFs. PhyB (Pfr) form can be made to revert to the PhyB (Pr) form by far-red light irradiation. (B) Red/far-red light-induced association and dissociation of PhyB-PIF. PIF-mEGFP fluorescence images are shown in HeLa cells expressing synPCB1.0, PhyB-mCherry-HRasCT, and PIF3-mEGFP upon the illumination with red and far-red light. Scale bar, 20 μm. (C) Membrane-recruited PIF-mEGFP was quantified by the fractional change in the fluorescence intensity of PIF-mEGFP at the cytoplasm under the condition of far-red light exposure. The average values (bold lines) are plotted as a function of time with the SD. The number of cells analyzed is n = 15.

To overcome this issue, we and another research group have developed a genetically encoded system for PCB synthesis in mammalian cells and fission yeasts (Figure 1) ^20,21^. In this system, PCB can be generated by coexpression of four genes in the mitochondria—heme oxygenase 1 (HO1), PCB:ferredoxin (Fd) oxidoreductase (PcyA), ferredoxin (Fd), and Fd-NADP+ reductase (Fnr)—all of which are derived from cyanobacteria (Figure 1A). Further increase in PCB synthesis has been achieved by knock-out (KO) or knock-down of the *biliverdin reductase A* (*BVRA*) gene, which catalyzes the generation of bilirubin and phycocyanorubin from biliverdin and PCB, respectively ^22^. The PCB synthesis system we have developed has been shown to work better than systems that externally add crude or even HPLC-purified PCBs ^23^, showing a repeated cycle of association and dissociation of PhyB-PIF on the second time scale (Figure 1B and 1C). However, a stable cell line that can sufficiently synthesize PCB and show an optogenetic response with the PhyB-PIF system in a genetically encoded manner has not yet been established. This is because the PCB synthesis system and stable expression methods have not been fully improved and optimized.

In this study, we present a further improvement of the PCB synthesis system in mammalian cells. As a result of several modifications, the latest PCB synthesis system, synPCB2.1, realizes two-fold greater PCB synthesis compared to the previous version. Additionally, we show that the combination of the Tol2 transposon system and doxycycline (dox)-inducible gene expression enables us to induce a high level of PCB synthesis in stable cell lines. These technical advances provide a versatile platform for the application of phytochrome-based optogenetics in a broader range of living organisms.

## Results and Discussion

### Remodeling ferredoxin and ferredoxin-NADP+ reductase improves PCB synthesis in mammalian cells

For the synthesis of PCB in cultured mammalian cells, it is necessary to express the above-named four genes in the mitochondria: *HO1*, *PcyA*, *Fd*, and *Fnr* ^20,21^(Figure 1A). In our system, the *HO1* and *PcyA genes* were originally derived from the thermophilic cyanobacterium *Thermosynechococcus* elongatus BP-1, while *Fd* and *Fnr* were obtained from the freshwater cyanobacterium *Synechocystis* sp. PCC6803. We first replaced *Fd* and *Fnr* from *Synechocystis* sp. PCC6803 with those from *Thermosynechococcus* elongatus BP-1. Second, we deleted the CpcD-like domain at the N-terminus of Fnr (truncated Fnr, tFnr). Fnr contains a CpcD-like domain at the N-terminus responsible for Fnr binding to phycobilisomes (Figure 2A) ^24^.

**Figure 2.**
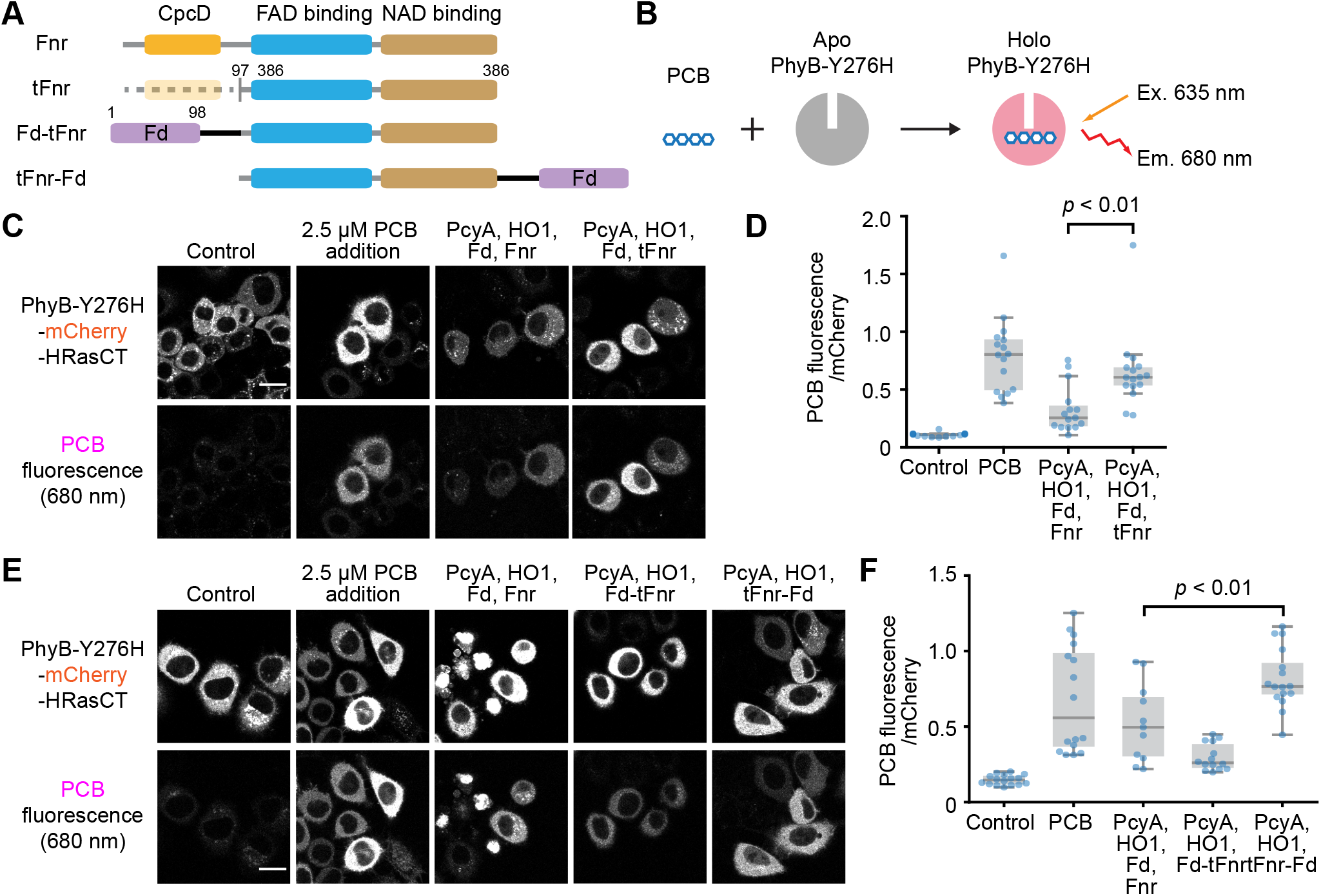
Improvement of PCB synthesis. (A) A schematic representation of Fnr and its chimeric proteins. tFnr, truncated Fnr. (B) PCB was measured by the fluorescence emitted from PCB binding to the PhyB-Y276H mutant. (C and E) Representative fluorescence images of PhyB-Y276H-mCherry-HRasCT (Upper) and PCB bound to PhyB-Y276H (Lower) under the indicated condition in HeLa cells. Scale bars, 20 μm. (D and F) Quantification of PCB synthesis. The fluorescence of PCB bound to PhyB-Y276H was divided by mCherry fluorescence, and shown as a box plot, in which the box extends from the first to the third quartile, with the whiskers denoting 1.5 times the interquartile range. Blue dots indicate individual cells. The numbers of cells in D are as follows: Control n = 16; PCB n = 16; PcyA, HO1, Fd, Fnr n = 15; PcyA, HO1, Fd, tFnr = 16. The numbers of cells in F are as follows: Control n = 16; PCB n = 16; PcyA, HO1, Fd, Fnr n = 11; PcyA, HO1, Fd-tFnr = 14; PcyA, HO1, tFnr-Fd = 16. *p*-values were obtained by unpaired two-tailed Welch’s unequal variance *t*-test.

We confirmed the capability of PCB synthesis by infra-red fluorescence emitted from PCB covalently bound to the PhyB-Y276H mutant (Figure 2B) ^25^. As a control, HeLa cells expressing PhyB-Y276H-Venus were treated with mock or purified 2.5 μM PCB, and it was confirmed that clear infra-red fluorescence was only emitted from only PCB-treated cells (Figure 2C). PCB production was observed by the transient expression of PcyA, HO1, Fd, and Fnr, all of which were derived from thermophilic cyanobacteria, and this was further increased by substitution of Fnr for tFnr (Figure 2C and 2D). This increase could be due to the increase in the expression level of tFnr by truncating the N-terminus. Of note, the mitochondrial targeting sequence (MTS) derived from human cytochrome C oxidase subunit VIII was fused to the N-terminus in all proteins to load these proteins into the mitochondrial matrix. The PCB fluorescence was normalized by dividing by the fluorescence intensity from PhyB-Y276H-mCherry mutants (Figure 2D) or PhyB-Y276H-mVenus mutants (Figure 3)

**Figure 3.**
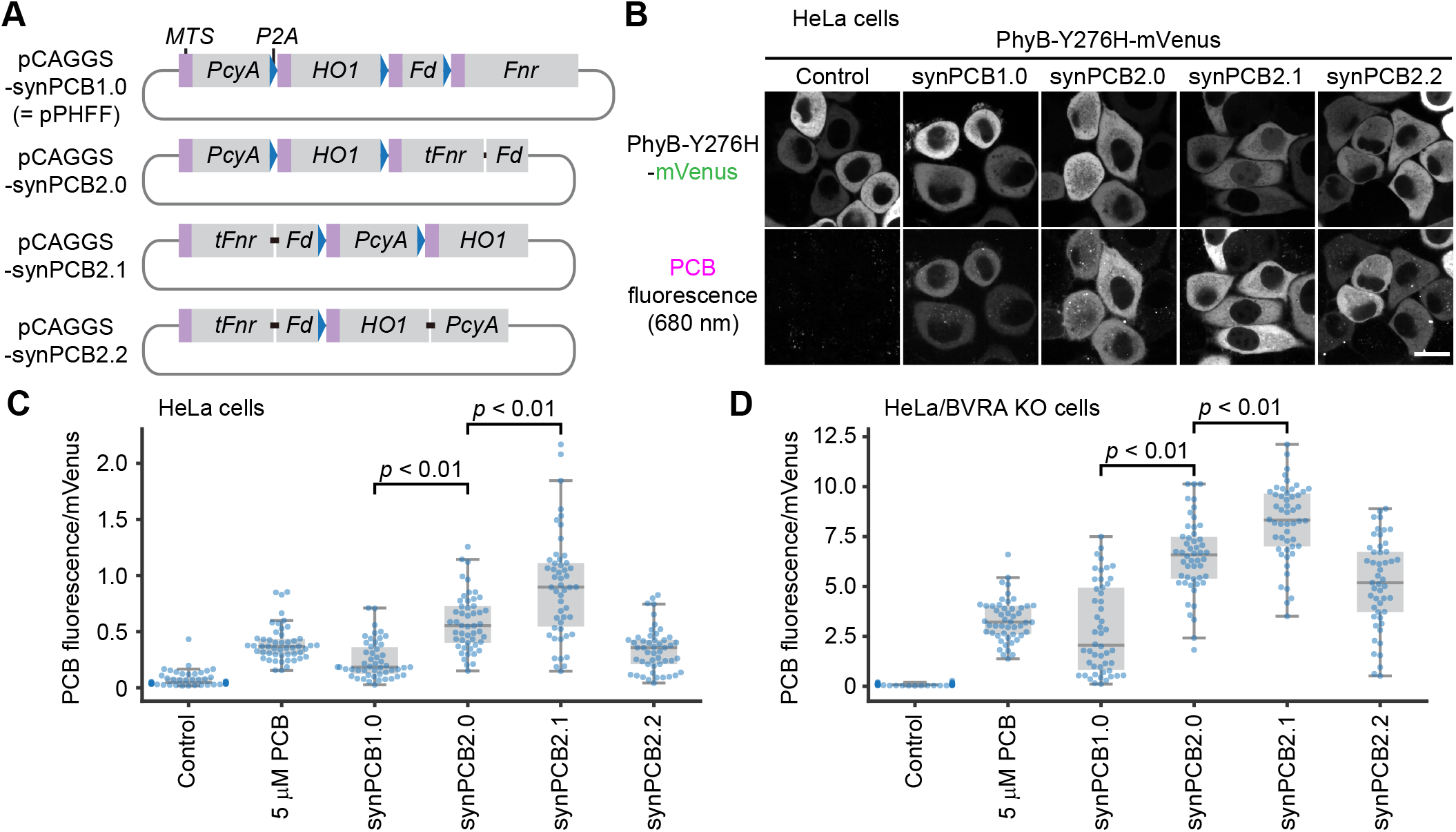
Polycistronic vectors for the synthesis of PCB. (A) Structure of the pCAGGS-synPCB plasmids. *MTS* and *P2A* encode the mitochondria-targeting sequence and self-cleaving 2A peptide. (B) Representative fluorescence images of PhyB-Y276H-mVenus (Upper) and PCB bound to PhyB-Y276H (Lower) under the indicated condition in HeLa cells. Scale bar, 10 μm. (C and D) PCB synthesis was quantified as in Figure 2D and 2F in HeLa cells (C) and HeLa/BVRA KO cells (D). The numbers of cells in C and D are n = 50 under all conditions. *p*-values were obtained by unpaired two-tailed Welch’s unequal variance *t*-test.

To achieve a greater increase in PCB synthesis, we focused on enzyme clustering or colocalization, which has been shown to enhance metabolic flux through high local concentrations of enzymes and intermediate metabolites ^26,27^. We have noticed that the expression of Fd is a limiting factor for PCB synthesis. In addition, Fd is known to form a heterodimer with Fnr ^28^, prompting us to generate a chimeric fusion protein to achieve higher activity of the Fd protein. Two chimeric proteins, Fd-tFnr and tFnr-Fd, were generated (Figure 2A), and were expressed with the PhyB-Y276H mutant, PcyA, and HO1 in HeLa cells (Figure 2E). Surprisingly, both chimeric proteins were competent to synthesize PCB, and tFnr-Fd showed higher PCB production than when Fd and full-length Fnr expressed separately (Figure 2E and 2F).

### synPCB: Polycistronic vectors for the synthesis of PCB

To easily introduce the genes required for PCB synthesis, we have developed a quad-cistronic vector for separate and equimolar expression of PcyA, HO1, Fd, and Fnr proteins, *i.e.*, pPHFF (Figure 3A) ^20^. Here, we made several modifications to Fd and Fnr, and to avoid confusion, we changed the name of the polycistronic vector for PCB synthesis from pPHFF to synPCB1.0. In synPCB1.0, the cDNAs of MTS-PcyA, MTS-HO1, MTS-Fd, and MTS-Fnr were concatenated with the cDNAs of a self-cleaving P2A peptide, enabling the polycistronic and stoichiometric expression of multiple proteins flanking the P2A peptide ^29^.

First, MTS-Fd-P2A-MTS-Fnr in the original SynPCB1.0 was replaced with MTS-tFnr-Fd to generate synPCB2.0 (Figure 3A and Figure S1). As in Figure 2, the expression of synPCB2.0 in HeLa cells expressing PhyB-Y276H showed an approximately 1.5-fold increase in PCB synthesis compared to the expression of the original SynPCB1.0 (Figure 3B and 3C). Next, we changed the order of the gene cassettes, namely, MTS-tFnr-Fd, MTS-PcyA, and MTS-HO1, and created synPCB2.1 (Figure 3A and Figure S1). The expression of synPCB2.1 further increased the activity of PCB synthesis by 3.9 times relative to the original SynPCB1.0 (Figure 3B and 3C). This result is consistent with the fact that Fd activity is the limiting factor, and the expression level decreased as it was moved towards the end of the 2A peptide-connected construct ^30^. Moreover, we generated a chimeric of HO1 and PcyA, because fusing these enzymes resulted in a huge increase in PCB production in *Bacillus subtilis* ^31^, and included it into the polycistronic vector, yielding synPCB2.2 (Figure 3A). However, unexpectedly, PCB production with synPCB2.2 was lower than that with synPCB2.0 and synPCB2.1 (Figure 3B and 3C). This may have been due to the steric hindrance between the HO1-PcyA and tFnr-Fd chimeric proteins. A similar trend was observed in HeLa cells depleted of BVRA (Figure 3D).

### Stable and drug-inducible PCB synthesis with transposon-mediated gene transfer

Finally, we attempted to establish stable cell lines for fully genetically encoded phytochrome-based optogenetics with the synPCB system. For this purpose, we used BVRA KO HeLa cells (HeLa/BVRA KO) for higher PCB production than in parental HeLa cells ^20^. First, we introduced reverse tetracycline-controlled transactivator (rtTA) with lentivirus into HeLa/BVRA KO cells (Figure 4A, left). The rtTA binds to tetracycline-responsive elements (TREs) in a doxycycline-dependent manner, and induces gene expression (Figure 4B).

**Figure 4.**
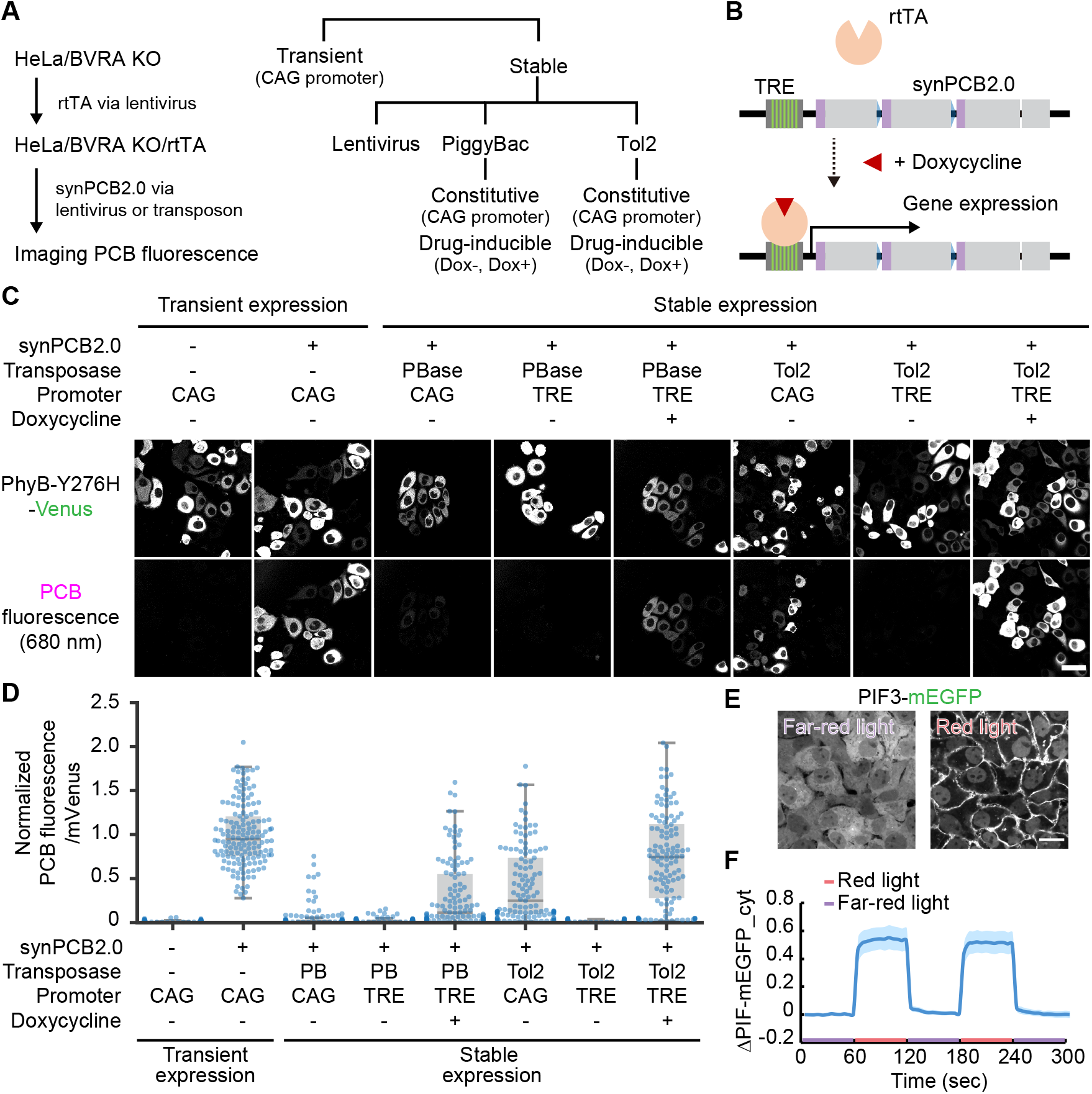
Stable and drug-inducible PCB synthesis with transposon-mediated gene transfer. (A and B) A scheme for the establishment of stable cell lines (A) and drug-inducible gene expression (B). (C) Representative fluorescence images of PhyB-Y276H-mVenus (Upper) and PCB bound to PhyB-Y276H (Lower) under the indicated condition in HeLa/BVRA KO/rtTA cells. CAG and TRE denote the CAGGS promoter and tetracycline-responsive element, respectively. Scale bar, 50 μm. (D) Quantification of PCB synthesis. The fluorescence of PCB fluorescence bound to PhyB-Y276H was divided by mVenus fluorescence, followed by normalization to the average PCB-bound PhyB-Y276H/mVenus value of 5 μM PCB-treated cells. The box extends from the first to the third quartile, with the whiskers denoting 1.5 times the interquartile range. Blue dots indicate individual cells. The numbers of cells in D are as follows: Transient, CAG n = 152; Transient, synPCB2.0 n = 150; PBase, CAG n = 121; PBase, TRE, Dox-n = 128; PBase, TRE, Dox+ n = 124; Tol2, CAG n = 121; Tol2, TRE, Dox-n = 120; Tol2, TRE, Dox+ n = 120. (E) Membrane translocation of PIF3-mEGFP was observed in HeLa/BVRA KO/rtTA/TRE-synPCB2.0/PhyB621-mCherry-HRasCT/PIF3-mEGFP cells treated with 1 μg/mL doxycycline for 1 day. Scale bar, 20 μm. (F) Membrane-recruited PIF-mEGFP was quantified as in Figure 1C. The average values (bold lines) are plotted as a function of time with the SD. The number of cells analyzed is n = 20.

We tested the lentivirus- and transposon-mediated gene transfer for constitutive or drug-inducible expression of synPCB2.0 (Figure 4A, right). The lentivirus-mediated gene transfer failed to synthesize PCB because of an unknown gene recombination. Therefore, we employed PiggyBac and Tol2 transposon-mediated gene transfer systems ^32,33^. As a control, CAG promoter-driven transient expression of synPCB2.0 with lipofection showed reasonably high PCB production (Figure 4C, second column). Meanwhile, the constitutive expression of synPCB2.0 through PiggyBac and Tol2 transposon showed smaller PCB fluorescence (Figure 4C, third and sixth columns), mainly because of the difference of the copy number between transient and stable expression. For the drug-inducible system, no PCB was synthesized without doxycycline treatment (Figure 4C, fourth and seventh columns). In both cell lines established with PiggyBac and Tol2, the cells treated with doxycycline exhibited higher PCB fluorescence than the doxycycline-untreated cells and constitutively expressing cells (Figure 4C, fifth and eighth columns). Among them, the drug-inducible expression of synPCB2.0 introduced with Tol2 transposon showed almost the same levels of PCB production as in the transient expression of synPCB2.0 (Figure 4D). An increase in PCB production could be achieved by the effect of rtTA, in which the transcriptional activity of the drug-inducible system is known to be higher than that of constitutive promoters ^34,35^. Several single-cell clones were obtained from cells with the drug-inducible system established by Tol2 transposon-mediated gene transfer (HeLa/BVRA KO/rtTA/TRE-synPCB2.0), and tested in regard to the doxycycline dose-response and time-course for PCB production; the optimal conditions were found to be 100-300 ng/mL doxycycline treatment for 20-40 hours (Figure S2).

To provide proof-of-concept, we examined whether the stable cell lines suffice to show red/far-red light-dependent PhyB-PIF binding. PhyB-mCherry-HRas C-terminus (HRasCT) and PIF3-mEGFP were additionally introduced into HeLa/BVRA KO/rtTA/TRE-synPCB2.0 cells through lentivirus, followed by single-cell cloning. As we expected, the cells treated with doxycycline uniformly demonstrated clear binding of PIF3-mEGFP and PhyB-mCherry-HRasCT, which was manifested by the membrane translocation retention of PIF3-mEGFP upon red light illumination, respectively (Figure 4E and 4F, Movie S1). This binding was restored repeatedly by far-red light illumination, indicating cytoplasmic retention of PIF3-mEGFP (Figure 4E and 4F, Movie S1).

In this study, we succeeded in improving the PCB synthesis system and establishing stable cell clones using the improved synPCB system. Our results will make it easier to apply PhyB- and cyanobacterial phytochrome (Cph)-based optogenetics in a fully genetically encoded manner ^36^. There were two key points in our improvement of PCB synthesis: the chimera of Fd and Fnr and the optimization of the order of the genes connected with the P2A sequence. The former enhances the activity of metabolic flux in PCB synthesis, while the latter optimizes the stoichiometry of these four factors. In addition, the PCB synthesis was further potentiated by introducing the doxycycline-inducible expression system. Such drug inducible systems allow not only the synthesis of PCB with appropriate timing but also a decrease in toxicity due to synPCB expression. Although the toxicity has not been observed in cultured cells (HeLa cells, mouse embryonic fibroblasts, and mouse embryonic stem cells) or fission yeast, the cytotoxicity due to synPCB expression should be minimized. These findings will open the door to the application of phytochrome-based optogenetics to higher organisms such as mice.

## Methods

### Plasmids

The cDNAs of *PcyA* and *HO1* were originally derived from *Thermosynechococcus* elongatus BP-1 ^20^. The cDNAs of *Fd* and *Fnr* of *Thermosynechococcus* elongatus BP-1 were synthesized with codon-optimization for the human genome by FASMAC (Kanagawa, Japan). The mitochondrial targeting sequence (MTS; MSVLTPLLLRGLTGSARRLP) was derived from human cytochrome C oxidase subunit VIII. The cDNAs were subcloned into vectors through conventional ligation with Ligation high Ver.2 (Toyobo, OSAKA, Japan) or NEBuilder HiFi DNA Assembly (New England Biolabs, Ipswich, MA) according to the manufacturer’s instructions. All plasmids used in this study are listed in Supplementary Table S1. The amino acid sequences of SynPCB2 and SynPCB3 are included in Figure S1.

### Reagents

PCB was purchased from Santa Cruz Biotechnology, dissolved in DMSO (5 mM), and stored at −30 °C. Doxycycline was obtained from Tokyo Chemical Industry (Tokyo; D45115), dissolved in DMSO (10 mg/mL), and stored at −30 °C.

### Cells

HeLa cells were purchased from the Human Science Research Resources Bank and maintained in DMEM (Nacalai tesque, Kyoto, Japan) supplemented with 10% fetal bovine serum (Sigma-Aldrich, St. Louis, MO). BVRA-KO HeLa cells (HeLa/BVRA KO) were established previously^20^. HEK-293T cells were obtained from Invitrogen (Carlsbad, CA) as Lenti-X 293 cells for lentivirus production. The HEK-293T cells were maintained in DMEM supplemented with 10% FBS.

### Establishment of stable cell lines

For transposon-mediated gene transfer, cells were transfected with the PiggyBac or Tol2 transposase expression vector (mPBase or pCAGGS-T2TP; Table S1) and the donor vectors using 293fectin (Thermo Fisher Scientific, Waltham, MA). One day after tge transfections, cells were selected by 10 μg/mL blasticidin S (InvivoGen, San Diego, CA) or 1.0 μg/mL puromycin (InvivoGen). For lentivirus-mediated gene transfer, HEK-293T cells were cotransfected with each pCSII vector, psPax2 (a gift from Didier Trono; Addgene plasmid # 12260), and pCMV-VSV-G-RSV-Rev (a gift from Dr. Miyoshi, RIKEN BioResource Center, Ibaraki, Japan), by using polyethyleneimine “Max” MW 40,000 (Polyscience, Warrington, PA). Virus-containing media were collected 48 hours after transfection, filtered, and applied to target cells with 10 μg/mL polybrene (Nacalai Tesque). Two days after infection, the infected cells were selected with the following antibiotics: 200 μg/mL hygromycin (Wako, Osaka, Japan), 800 μg/mL G418 (InvivoGen), 10 μg/mL blasticidin S (InvivoGen), 1.0 μg/mL puromycin (InvivoGen), and 10 μg/mL zeocin (InvivoGen). Single-cell clones were isolated by the limiting dilution method.

### Fluorescence imaging

Fluorescence imaging was conducted essentially as previously reported ^20^. Briefly, HeLa cells were plated on poly–L-lysine (PLL)-coated 35-mm glass-base dishes. For transient expression, the plasmids were transfected by using 293fectin (Invitrogen) according to the following; PhyB:PIF-mEGFP:synPCB1.0 = 50:1:50 (Figure 1); PhyB-Y276H:PcyA:HO1:Fd:Fnr = 2:1:1:1:1 (Figure 2C and 2D); PhyB-Y276H:PcyA:HO1:Fd-Fnr chimera = 2:1:1:2 (Figure 2E and 2F); PhyB-Y276H:synPCBs = 1:1 (Figure 3). After 48 h, the medium was replaced with FluoroBrite D-MEM (Thermo Fisher Scientific) supplemented with 1% GlutaMAX (Thermo Fisher Scientific) and 0.1% BSA. PCB was added and culture media was incubated for 30 min, followed by a medium change.

For confocal fluorescence imaging, cells were imaged with a TCS SP5 microscope (Leica Microsystems, Mannheim, Germany) or IX83 inverted microscope (Olympus, Tokyo) equipped with an sCMOS camera (Prime; Photometrics, Tucson, AZ), and a spinning disk confocal unit (CSU-W1; Yokogawa Electric Corporation, Tokyo) illuminated with a laser merge module containing 488 nm, 561 nm, and 640 nm lasers. In the Leica TSC SP5 confocal imaging system, an oil immersion objective lens (HCX PL APO 63×1.4–0.6 oil; Leica Microsystems) was used. The excitation laser and fluorescence filter settings were as follows: excitation laser, 488 nm (mEGFP or mVenus), 543 nm (mCherry), and 633 nm (PCB fluorescence); excitation dichroic mirror, TD 488/543/633 dichroic mirror; detector, HyD 520–590 nm (mEGFP), HyD 600-650 nm (mCherry), and HyD 670–720 nm (PCB fluorescence). In the Olympus imaging system, an oil immersion objective lens (UPLANSAPO 60X, N.A. 1.35; Olympus) or an air/dry objective lens (UPLFLN40X, N.A. 0.75; Olympus) was used. The excitation laser and fluorescence filter settings were as follows: excitation laser, 488 nm (mEGFP or mVenus), 561 nm (mCherry), and 640 nm (PCB fluorescence); excitation dichroic mirror, DM 405/488/561 dichroic mirror, 500–550 nm (mEGFP), 580–654 nm (mCherry), and 665–705 nm (PCB fluorescence). For red and far-red light illumination for the binding and dissociation between PhyB and PID3, LEDs for red (625 nm) and far-red (735 nm) light were purchased from Optocode and controlled manually.

## Associated content

The Supporting Information is available free of charge on the ACS Publications website at the following DOI:

Figure S1: The amino acid sequence of synPCB vectors.

Figure S2: Dose-response and time-course of doxycycline-induced PCB synthesis.

Table S1: Plasmids used in this study.

Movie S1: Repeated translocation of PIF-mEGFP to the plasma membrane in HeLa cells.

## Notes

The authors declare no competing financial interest.

## Acknowledgments

We thank Emi Ebine and Kaori Onoda for their assistance and all members of the Aoki Laboratory for helpful discussions. Plasmid DNAs were provided by K. Yusa and A. Bradley (piggyBAC), K. Kawakami (Tol2), J. Miyazaki (pCAGGS), H. Miyoshi (pCSII and pCMV-VSV-GRSV-Rev), Dr. D. Trono (psPAX2), and Dr. F. Zhang (lentiCRISPR v2). This research was supported by CREST, JST (JPMJCR1654), and JSPS KAKENHI grants (nos. 16KT0069, 16H01425 “Resonance Bio”, 18H04754 “Resonance Bio”, 18H02444, 19H05798, no.19K16050).

## Abbreviations

PhyB: phytochrome B
PIF: phytochrome-interacting factor
PCB: phycocyanobilin
PcyA: phycocyanobilin:ferredoxin oxidoreductase
HO1: heme oxygenase
Fd: ferredoxin
FNR: ferredoxin-NADP+-reductase
BV: biliverdin IX-alpha
BVRA: biliverdin reductase A

## Supplementary Information

**Figure S1.**
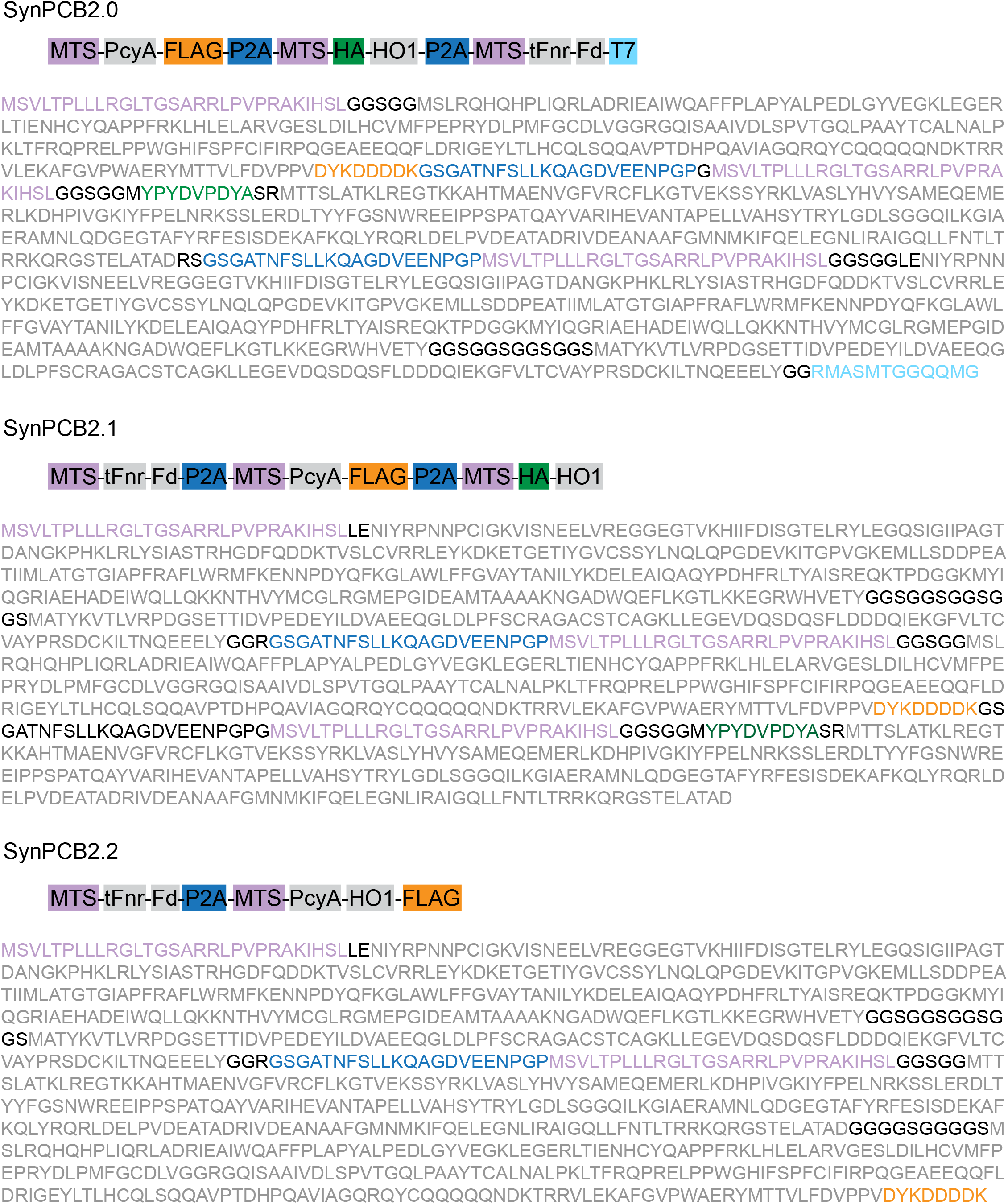
The amino acid sequence of synPCB vectors. The amino acid sequence of each synPCB vector is shown with the indicated color.

**Figure S2.**
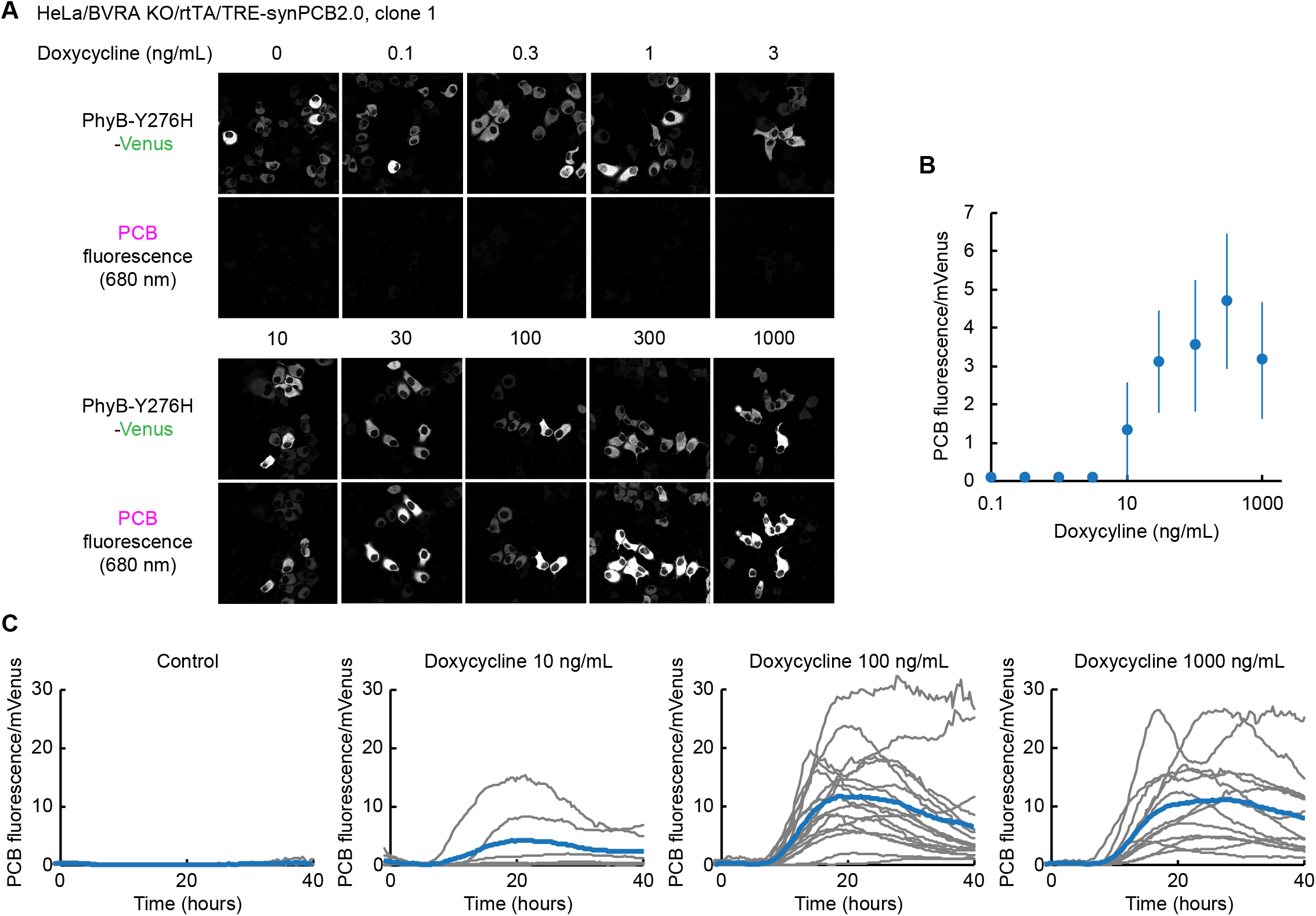
Dose-response and time-course curves of doxycycline-induced PCB synthesis. (A) HeLa/BVRA KO/rtTA/TRE-synPCB2.0 cells were transfected with the PhyB-Y276H-mVenus expression plasmid. One day after transfection, the cells were treated with the indicated concentration of doxycycline. One day after doxycycline treatment, the fluorescence of PhyB-Y276H-mVenus (Upper) and the PCB fluorescence bound to PhyB-Y276H (Lower) were imaged. (B) Quantification of PCB synthesis in A. The fluorescence of PCB bound to PhyB-Y276H was divided by mVenus fluorescence. The average PCB fluorescence/mVenus fluorescence values are plotted with the SD. The number of cells is more than 30 cells under all conditions. (C) HeLa/BVRA KO/rtTA/TRE-synPCB2.0 cells were transfected with PhyB-Y276H-mVenus expression plasmid. One day after transfection, the cells were treated with the indicated concentration of doxycycline, and time-lapse imaging started. The time-courses of PCB fluorescence/mVenus fluorescence values are plotted. Gray and blue lines represent individual cells and averaged data, respectively. The numbers of cells were as follows: Control n = 4; Doxycycline 10 ng/mL n = 7; Doxycycline 100 ng/mL n = 17; Doxycycline 1000 ng/mL n = 13.

**Table S1.**
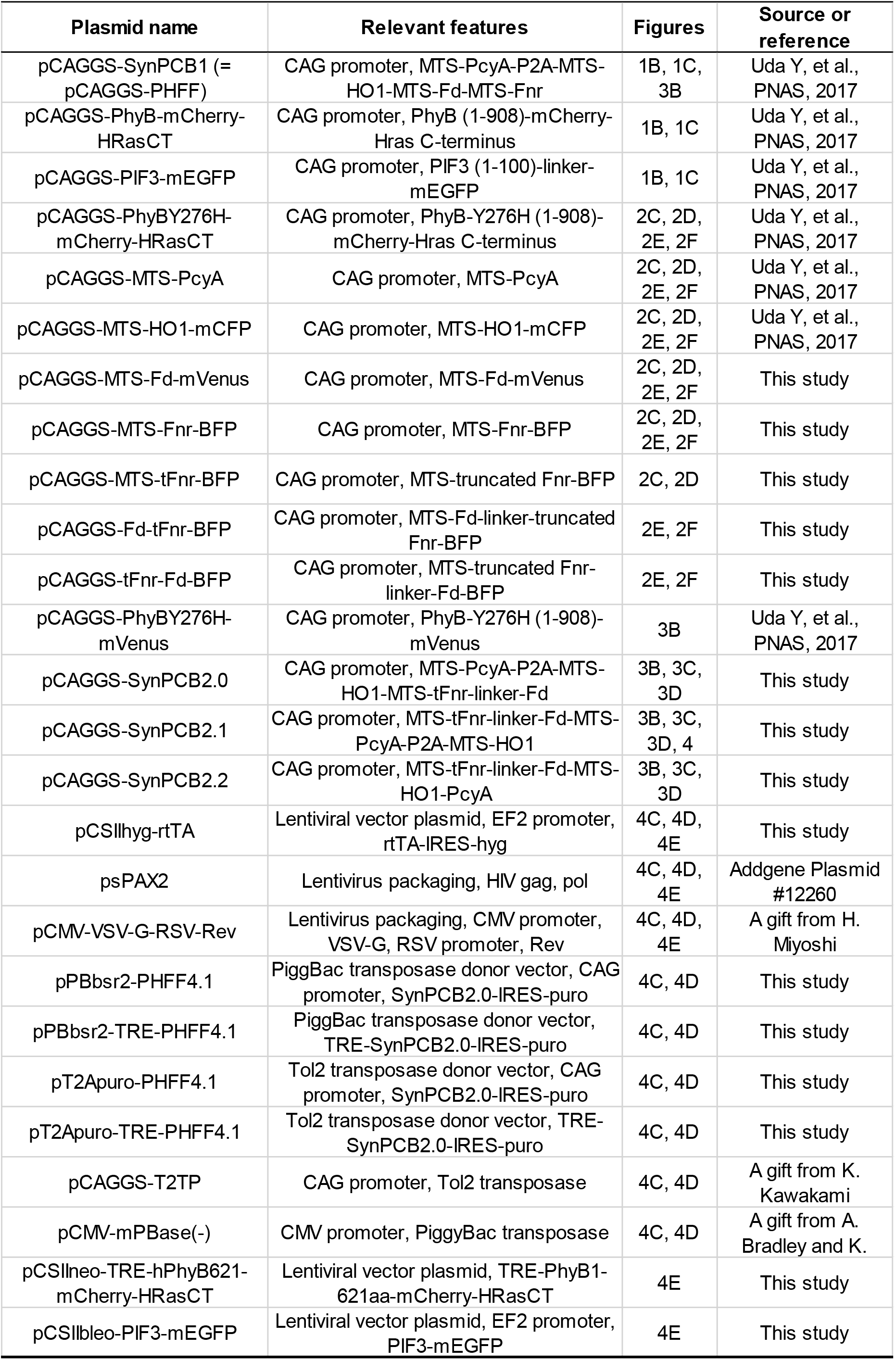
Plasmids used in this study.

| Plasmid name | Relevant features | Figures | Source or reference |
| --- | --- | --- | --- |
| pCAGGS-SynPCB1 (= pCAGGS-PHFF) | CAG promoter, MTS-PcyA-P2A-MTS-HO1-MTS-Fd-MTS-Fnr | 1B, 1C, 3B | Uda Y, et al., PNAS, 2017 |
| pCAGGS-PhyB-mCherry-HRasCT | CAG promoter, PhyB (1-908)-mCherry-Hras C-terminus | 1B, 1C | Uda Y, et al., PNAS, 2017 |
| pCAGGS-PIF3-mEGFP | CAG promoter, PIF3 (1-100)-linker-mEGFP | 1B, 1C | Uda Y, et al., PNAS, 2017 |
| pCAGGS-PhyBY276H-mCherry-HRasCT | CAG promoter, PhyB-Y276H (1-908)-mCherry-Hras C-terminus | 2C, 2D, 2E, 2F | Uda Y, et al., PNAS, 2017 |
| pCAGGS-MTS-PcyA | CAG promoter, MTS-PcyA | 2C, 2D, 2E, 2F | Uda Y, et al., PNAS, 2017 |
| pCAGGS-MTS-HO1-mCFP | CAG promoter, MTS-HO1-mCFP | 2C, 2D, 2E, 2F | Uda Y, et al., PNAS, 2017 |
| pCAGGS-MTS-Fd-mVenus | CAG promoter, MTS-Fd-mVenus | 2C, 2D, 2E, 2F | This study |
| pCAGGS-MTS-Fnr-BFP | CAG promoter, MTS-Fnr-BFP | 2C, 2D, 2E, 2F | This study |
| pCAGGS-MTS-tFnr-BFP | CAG promoter, MTS-truncated Fnr-BFP | 2C, 2D | This study |
| pCAGGS-Fd-tFnr-BFP | CAG promoter, MTS-Fd-linker-truncated Fnr-BFP | 2E, 2F | This study |
| pCAGGS-tFnr-Fd-BFP | CAG promoter, MTS-truncated Fnr-linker-Fd-BFP | 2E, 2F | This study |
| pCAGGS-PhyBY276H-mVenus | CAG promoter, PhyB-Y276H (1-908)-mVenus | 3B | Uda Y, et al., PNAS, 2017 |
| pCAGGS-SynPCB2.0 | CAG promoter, MTS-PcyA-P2A-MTS-HO1-MTS-tFnr-linker-Fd | 3B, 3C, 3D | This study |
| pCAGGS-SynPCB2.1 | CAG promoter, MTS-tFnr-linker-Fd-MTS-PcyA-P2A-MTS-HO1 | 3B, 3C, 3D, 4 | This study |
| pCAGGS-SynPCB2.2 | CAG promoter, MTS-tFnr-linker-Fd-MTS-HO1-PcyA | 3B, 3C, 3D | This study |
| pCSIIhyg-rtTA | Lentiviral vector plasmid, EF2 promoter, rtTA-IRES-hyg | 4C, 4D, 4E | This study |
| psPAX2 | Lentivirus packaging, HIV gag, pol | 4C, 4D, 4E | Addgene Plasmid #12260 |
| pCMV-VSV-G-RSV-Rev | Lentivirus packaging, CMV promoter, VSV-G, RSV promoter, Rev | 4C, 4D, 4E | A gift from H. Miyoshi |
| pPBbsr2-PHFF4.1 | PiggBac transposase donor vector, CAG promoter, SynPCB2.0-IRES-puro | 4C, 4D | This study |
| pPBbsr2-TRE-PHFF4.1 | PiggBac transposase donor vector, TRE-SynPCB2.0-IRES-puro | 4C, 4D | This study |
| pT2Apuro-PHFF4.1 | Tol2 transposase donor vector, CAG promoter, SynPCB2.0-IRES-puro | 4C, 4D | This study |
| pT2Apuro-TRE-PHFF4.1 | Tol2 transposase donor vector, TRE-SynPCB2.0-IRES-puro | 4C, 4D | This study |
| pCAGGS-T2TP | CAG promoter, Tol2 transposase | 4C, 4D | A gift from K. Kawakami |
| pCMV-mPBase(-) | CMV promoter, PiggyBac transposase | 4C, 4D | A gift from A. Bradley and K. |
| pCSIIneo-TRE-hPhyB621-mCherry-HRasCT | Lentiviral vector plasmid, TRE-PhyB1-621aa-mCherry-HRasCT | 4E | This study |
| pCSIIbleo-PIF3-mEGFP | Lentiviral vector plasmid, EF2 promoter, PIF3-mEGFP | 4E | This study |

**Movie S1. Repeated translocation of PIF-mEGFP to the plasma membrane in HeLa cells**. PIF3-mEGFP localization was controlled by red and far-red light illumination in HeLa/BVRA KO cells stably expressing rtTA, synPCB2.0, PhyB621-mCherry-HRasCT, and PIF3-mEGFP. Timestamp shows time in minutes:seconds.

## References

(1) Kolar, K.; Knobloch, C.; Stork, H.; Žnidarič, M.; Weber, W. OptoBase: A Web Platform for Molecular Optogenetics. ACS Synth. Biol. 2018, 7 (7), 1825–1828.

(2) Spiltoir, J. I.; Tucker, C. L. Photodimerization Systems for Regulating Protein-Protein Interactions with Light. Curr. Opin. Struct. Biol. 2019, 57, 1–8.

(3) Goglia, A. G.; Toettcher, J. E. A Bright Future: Optogenetics to Dissect the Spatiotemporal Control of Cell Behavior. Curr. Opin. Chem. Biol. 2019, 48, 106–113.

(4) Aoki, K.; Kumagai, Y.; Sakurai, A.; Komatsu, N.; Fujita, Y.; Shionyu, C.; Matsuda, M. Stochastic ERK Activation Induced by Noise and Cell-to-Cell Propagation Regulates Cell Density-Dependent Proliferation. Mol. Cell 2013, 52 (4), 529–540.

(5) Aoki, K.; Kondo, Y.; Naoki, H.; Hiratsuka, T.; Itoh, R. E.; Matsuda, M. Propagating Wave of ERK Activation Orients Collective Cell Migration. Dev. Cell 2017, 43 (3), 305–317.e5.

(6) Katsura, Y.; Kubota, H.; Kunida, K.; Kanno, A.; Kuroda, S.; Ozawa, T. An Optogenetic System for Interrogating the Temporal Dynamics of Akt. Sci. Rep. 2015, 5, 14589.

(7) Idevall-Hagren, O.; Dickson, E. J.; Hille, B.; Toomre, D. K.; De Camilli, P. Optogenetic Control of Phosphoinositide Metabolism. Proc. Natl. Acad. Sci. U. S. A. 2012, 109 (35), E2316–E2323.

(8) Konermann, S.; Brigham, M. D.; Trevino, A.; Hsu, P. D.; Heidenreich, M.; Cong, L.; Platt, R. J.; Scott, D. A.; Church, G. M.; Zhang, F. Optical Control of Mammalian Endogenous Transcription and Epigenetic States. Nature 2013, 500 (7463), 472–476.

(9) Duan, L.; Che, D.; Zhang, K.; Ong, Q.; Guo, S.; Cui, B. Optogenetic Control of Molecular Motors and Organelle Distributions in Cells. Chem. Biol. 2015, 22 (5), 671–682.

(10) Kennedy, M. J.; Hughes, R. M.; Peteya, L. A.; Schwartz, J. W.; Ehlers, M. D.; Tucker, C. L. Rapid Blue-Light-Mediated Induction of Protein Interactions in Living Cells. Nat. Methods 2010, 7 (12), 973–975.

(11) Guntas, G.; Hallett, R. A.; Zimmerman, S. P.; Williams, T.; Yumerefendi, H.; Bear, J. E.; Kuhlman, B. Engineering an Improved Light-Induced Dimer (iLID) for Controlling the Localization and Activity of Signaling Proteins. Proc. Natl. Acad. Sci. U. S. A. 2015, 112 (1), 112–117.

(12) Smith, H.; Whitelam, G. C. The Shade Avoidance Syndrome: Multiple Responses Mediated by Multiple Phytochromes. Plant Cell Environ. 1997, 20 (6), 840–844.

(13) Legris, M.; Ince, Y. Ç.; Fankhauser, C. Molecular Mechanisms Underlying Phytochrome-Controlled Morphogenesis in Plants. Nat. Commun. 2019, 10 (1), 5219.

(14) Shimizu-Sato, S.; Huq, E.; Tepperman, J. M.; Quail, P. H. A Light-Switchable Gene Promoter System. Nat. Biotechnol. 2002, 20 (10), 1041–1044.

(15) Levskaya, A.; Weiner, O. D.; Lim, W. A.; Voigt, C. A. Spatiotemporal Control of Cell Signalling Using a Light-Switchable Protein Interaction. Nature 2009, 461 (7266), 997–1001.

(16) Toettcher, J. E.; Weiner, O. D.; Lim, W. A. Using Optogenetics to Interrogate the Dynamic Control of Signal Transmission by the Ras/Erk Module. Cell 2013, 155 (6), 1422–1434.

(17) Kohchi, T.; Kataoka, H.; Linley, P. J. Biosynthesis of Chromophores for Phytochrome and Related Photoreceptors. Plant Biotechnol. 2005, 22 (5), 409–413.

(18) Mukougawa, K.; Kanamoto, H.; Kobayashi, T.; Yokota, A.; Kohchi, T. Metabolic Engineering to Produce Phytochromes with Phytochromobilin, Phycocyanobilin, or Phycoerythrobilin Chromophore in Escherichia Coli. FEBS Lett. 2006, 580 (5), 1333–1338.

(19) Pham, V. N.; Kathare, P. K.; Huq, E. Phytochromes and Phytochrome Interacting Factors. Plant Physiol. 2018, 176 (2), 1025–1038.

(20) Uda, Y.; Goto, Y.; Oda, S.; Kohchi, T.; Matsuda, M.; Aoki, K. Efficient Synthesis of Phycocyanobilin in Mammalian Cells for Optogenetic Control of Cell Signaling. Proceedings of the National Academy of Sciences 2017, 114 (45), 11962–11967.

(21) Kyriakakis, P.; Catanho, M.; Hoffner, N.; Thavarajah, W.; Hu, V. J.; Chao, S.-S.; Hsu, A.; Pham, V.; Naghavian, L.; Dozier, L. E.; et al. Biosynthesis of Orthogonal Molecules Using Ferredoxin and Ferredoxin-NADP+ Reductase Systems Enables Genetically Encoded PhyB Optogenetics. ACS Synth. Biol. 2018, 7 (2), 706–717.

(22) Kutty, R. K.; Maines, M. D. Purification and Characterization of Biliverdin Reductase from Rat Liver. J. Biol. Chem. 1981, 256 (8), 3956–3962.

(23) Oda, S.; Uda, Y.; Goto, Y.; Miura, H.; Aoki, K. CHAPTER 7:Optogenetic Tools for Quantitative Biology: The Genetically Encoded PhyB–PIF Light-Inducible Dimerization System and Its Application for Controlling Signal Transduction. In Optogenetics; 2018; pp 137–148.

(24) van Thor, J. J.; Gruters, O. W.; Matthijs, H. C.; Hellingwerf, K. J. Localization and Function of ferredoxin:NADP(+) Reductase Bound to the Phycobilisomes of Synechocystis. EMBO J. 1999, 18 (15), 4128–4136.

(25) Su, Y.-S.; Lagarias, J. C. Light-Independent Phytochrome Signaling Mediated by Dominant GAF Domain Tyrosine Mutants of Arabidopsis Phytochromes in Transgenic Plants. Plant Cell 2007, 19 (7), 2124–2139.

(26) Lee, H.; DeLoache, W. C.; Dueber, J. E. Spatial Organization of Enzymes for Metabolic Engineering. Metab. Eng. 2012, 14 (3), 242–251.

(27) Conrado, R. J.; Varner, J. D.; DeLisa, M. P. Engineering the Spatial Organization of Metabolic Enzymes: Mimicking Nature’s Synergy. Curr. Opin. Biotechnol. 2008, 19 (5), 492–499.

(28) Kurisu, G.; Kusunoki, M.; Katoh, E.; Yamazaki, T.; Teshima, K.; Onda, Y.; Kimata-Ariga, Y.; Hase, T. Structure of the Electron Transfer Complex between Ferredoxin and Ferredoxin-NADP(+) Reductase. Nat. Struct. Biol. 2001, 8 (2), 117–121.

(29) Kim, J. H.; Lee, S. R.; Li, L. H.; Park, H. J.; Park, J. H.; Lee, K. Y.; Kim, M. K.; Shin, B. A.; Choi, S. Y. High Cleavage Efficiency of a 2A Peptide Derived from Porcine Teschovirus-1 in Human Cell Lines, Zebrafish and Mice. PLoS One 2011, 6 (4), 1–8.

(30) Liu, Z.; Chen, O.; Wall, J. B. J.; Zheng, M.; Zhou, Y.; Wang, L.; Ruth Vaseghi, H.; Qian, L.; Liu, J. Systematic Comparison of 2A Peptides for Cloning Multi-Genes in a Polycistronic Vector. Sci. Rep. 2017, 7 (1), 2193.

(31) Castillo-Hair, S. M.; Baerman, E. A.; Fujita, M.; Igoshin, O. A.; Tabor, J. J. Optogenetic Control of Bacillus Subtilis Gene Expression. Nat. Commun. 2019, 10 (1), 3099.

(32) Yusa, K.; Rad, R.; Takeda, J.; Bradley, A. Generation of Transgene-Free Induced Pluripotent Mouse Stem Cells by the piggyBac Transposon. Nat. Methods 2009, 6 (5), 363–369.

(33) Kawakami, K.; Noda, T. Transposition of the Tol2 Element, an Ac-Like Element from the Japanese Medaka Fish Oryzias Latipes, in Mouse Embryonic Stem Cells. Genetics 2004, 166 (2), 895–899.

(34) Yin, D. X.; Zhu, L.; Schimke, R. T. Tetracycline-Controlled Gene Expression System Achieves High-Level and Quantitative Control of Gene Expression. Anal. Biochem. 1996, 235 (2), 195–201.

(35) Hioki, H.; Kuramoto, E.; Konno, M.; Kameda, H.; Takahashi, Y.; Nakano, T.; Nakamura, K. C.; Kaneko, T. High-Level Transgene Expression in Neurons by Lentivirus with Tet-Off System. Neurosci. Res. 2009, 63 (2), 149–154.

(36) Reichhart, E.; Ingles-Prieto, A.; Tichy, A. M.; McKenzie, C.; Janovjak, H. A Phytochrome Sensory Domain Permits Receptor Activation by Red Light. Angewandte Chemie - International Edition 2016, 55 (21), 6339–6342.

